# Influence of Strontium on the Physical, Mechanical and *In-Vitro* Bioactivity of Glass Ionomer Cements

**DOI:** 10.1101/870873

**Authors:** Yiyu Li

**Affiliations:** Department of General Dentistry, Peking University, Beijing, 100081

**Keywords:** GIC, Strontium, Compressive Strength, Flexural Strength, SBF, *In-vitro*

## Abstract

In this work, we investigated the effects of strontium incorporation in the glass phase of glass ionomer cements (GIC). Three different glass compositions were synthesized with 0, 5, and 10 mol% of SrO addition. GICs were prepared by the addition of 50 wt% polyacrylic acid (PAA) at powder to liquid ratio of 1:1.5. Initial characterization on the cement series was to study their rheological behavior. Cements represented working times between 50-64 seconds and setting times of 356-452 seconds. Rheological results indicated that the addition of strontium decreases the working and setting times of the cements. To analyze the mechanical properties, compressive and flexural strength studies were performed after 1, 10, and 30 days incubation in simulated body fluid. The compressive strength of the cements increased as a function of incubation time, with the strontium containing compositions showing the highest strength at 34 MPa and after 30 days of incubation. Biaxial flexural strength of the cements was not significantly affected by the composition and maturation time and ranged between 13.4 to 16.3 MPa. *In-vitro* bioactivity of the cements was analyzed using SBF trials and after 1, 10, and 30 days incubation periods. Strontium containing cements, showed higher solubility with higher amounts of calcium phosphate surface depositions only after 10 days incubation. The elemental identifications of the surface depositions indicated high amounts of Ca, P and Zn are present on the surface of SBF incubated samples.

## 1 Introduction

The development of dental adhesives for restorative dentistry originated with the work of Bouncier in the 1950s, in trials of bonding resin to etched enamel[1]. Since then, different compositions of chemically adhesive materials have been developed, with the introduction of zinc polycarboxylate cement (ZPC) in the 1960s and glass ionomer cements (GICs) following shortly after[2]. GICs were initially developed by Wilson and Kent as restoratives for dental applications in the 1970s[3]. The original GICs were water-based materials that set by an acid-base reaction. Further modifications of the chemical composition of these types of cements by Wilson and McLean, improved their physical properties[4]. Since their initial introduction, they have found versatile applications in the clinical dentistry such as linings, luting, and aesthetic restorations. GICs are hybrids of the polycarboxylate and silicate cements, with properties of both silicate cements such as translucency and fluoride release, and characteristics of polycarboxylate cements such chemical bond to tooth[5, 6]. GICs all contain a silicate-based glass phase and a polymer base in the form of an acid such as polyacrylic or tartaric acid and water. The first commercial GIC consisted of an aluminosilicate glass and an aqueous solution of polyacrylic acid[7].

Since the discovery of the GICs and their clinical use in dentistry, no significant adverse reaction has been reported[8]. In dental applications, the beneficial effects of GICs include adhesion to tooth mineral and release of fluoride ions that are thought to confer resistance against dental caries[9]. Also, freshly mixed GICs can chemically bond to both bone tissue and metal surfaces, which is beneficial in the sense that they do not rely solely on mechanical interaction to achieve fixation of a cement or implant to bone[10]. Conventional GICs adhere to untreated enamel and dentine, bone and base metals. Some GICs exhibit osteoconductive properties after implantation into bone[11]. Due to their superior biocompatibility, their capacity to adhere to surgical metals and chemically bond to the skeletal tissue, and also due to their unique setting reaction without heat generation and volumetric shrinkage, much attention has been focused on the development of GICs for use as bone cements[12, 13].

The setting reaction that occurs in all GICs is based on an acid-base reaction[12]. A hydrous polycarboxylic acid solution, conventionally PAA, reacts with an ion-releasing glass structure, which degrades to form a hydrogel or polysalt matrix[14]. The acid attacks the surface of the glass particles, which leads to surface degradation of the glass particles and release of metallic cations into the solution. The metallic cations, such as Ca^2+^, crosslink with the carboxylate groups of polyacrylate chains resulting in the formation of hard cement[15]. The resulting set cement consists of unreacted glass particles that have been embedded in a polysalt matrix[15].

The chemistry of the glass composition significantly affects the properties of the resultant cement composition[16]. The reactivity of the glass phase depends on the structure of the glass, the number of modifying cations, their valency and then on their acidity/basicity[17]. Inclusion of network modifiers such as Sr^2+^ can increase the number of non-bridging oxygens and make the glass network disrupted[18, 19]. This modified glass structure is susceptible to acid attack, which is an important property in degradable glasses. Strontium in the body behaves in a very similar fashion to Ca, where it is mostly accumulated in skeletal tissue[20, 21]. Research by Hao et al investigated the substitution of Ca by Sr into hydroxyapatite, as Sr is one of the divalent ions that can replace Ca[22]. Their study demonstrated that Sr can participate effectively in the re-mineralization process of the bone[22]. Another study by Tripathi et al have shown that the released Sr ions from glass structure can show anti-cariogenic properties[23]. The main goal of this study is to investigate how Sr incorporation can affect the properties of the GICs. In this study, we have incorporated SrO in the glass chemistry and evaluated the physical, mechanical and biological behavior of the resultant cements.

## 2 Materials & Methods

### 2.1 Glass Synthesis

To prepare the glass powders, three different glass compositions were synthesized; a silicate-based control glass, and two strontium-containing glass compositions. Detailed compositions of synthesized glass powders are presented in Table 1. To synthesize the glass, the traditional melt-quench method was employed. First, the analytical grades of reagents were mixed for 30 minutes and then oven-dried at 110°C for 2 hours. The glass batch was then melted in a platinum crucible at 1360°C for 4 hours. The molten glass was poured in cold water, and the resulting frits were dried overnight and pulverized to glass powders with particle sizes of ≤45*μ*m.

**Table 1.**
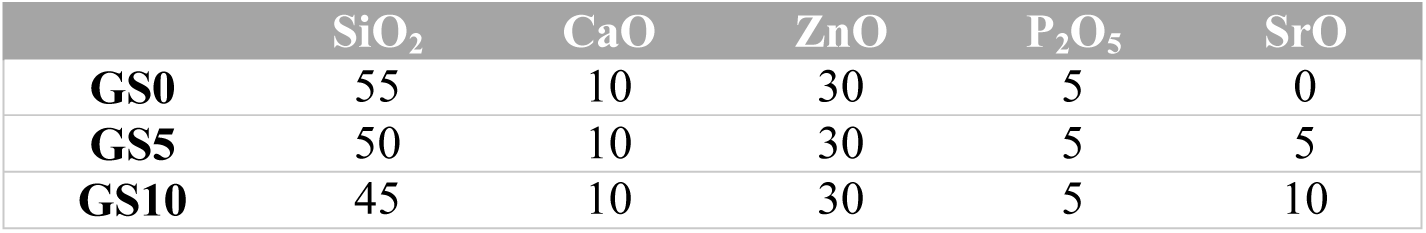
Glass compositions (Mol.%).

### 2.2 Glass Ionomer Cement Formulation

Glass ionomer cements were prepared by mixing the synthesized glass powders, polyacrylic acid (PAA) and deionized water on a glass slide. The formulation used (Table 2) for the cement preparation was at 50 wt.% PAA and powder to liquid ratio of 1:1.5.

**Table 2.** Cement formulation.

|  | Glass | PAA | Water |
| --- | --- | --- | --- |
| **P/L 1:1.5** | 1.00 g | 0.75 g | 0.75 ml |

### 2.3 Working/Setting Time Measurements

Working time was defined as the time from the start of the mixing of the cement until it is possible to manipulate the mix. The setting time of the cements was tested following ISO9917 standard which defines the standard for timing the setting times of glass ionomer cements. Briefly, cements were prepared and kept at 37°C during setting time analysis. Setting time was measured by lowering a 400 g mass attached to a needle into the cement disks. Setting time was defined as the time it takes for the needle to make a complete indent on the surface of cement. Each measurement was conducted on 5 samples to ensure reproducibility.

### 2.4 Compressive Strength

To evaluate the compressive strength first cements cylinders with dimensions of 15×7 mm were prepared. Once the cements were set, they were transferred to simulated body fluid and were kept for 1,10, and 30 days in an incubator at 37°C. At the end of each incubation time period, the cements were tested for their compressive strength in a Universal Testing Machine at a crosshead speed of 1 mm/min.

### 2.5 Biaxial Flexural Strength

To measure the flexural strength of the cement series, disks with dimensions of 3×10 mm were prepared. The set cement disks were kept in simulated body fluid for 1, 10, and 30 days prior to testing. The cement samples were tested on a biaxial flexural fixture, where the disks were loaded by a piston from above and supported by three balls from below. The measurements were performed on a Universal Testing Machine at a crosshead speed of 0.5 mm/min.

### 2.6 *In-Vitro* Bioactivity Testing

To study the *in-vitro* bioactivity of the cement samples, Simulated Body Fluid (SBF) solution was prepared following the formula and method proposed by Kokubo et al[24]. First cement disks with dimensions of 3×10 mm were prepared. Then each specimen was immersed in 10 ml of the SBF solution and stored in an incubator at 37°C for 1, 10, and 30 days. After the end of each incubation time period, specimens were removed from the solution and dried to be analyzed for the surface depositions with a scanning electron microscope (SEM).

### 2.7 Scanning Electron Microscopy

Imaging of the SBF incubated samples was performed using scanning electron microscope SEM (Philips 30XLFEG, Netherlands). Samples were gold coated for 30 seconds using a sputter coater

## 3 Results & Discussion

This study investigates the physical and biological effects of strontium incorporation in the glass composition of glass ionomer cements. Three different glass compositions with varying amount (0, 5 and 10 mol%) of SrO were prepared. The glass compositions are presented in Table 1. Glass frits were pulverized to glass powders with particle sizes of less than 45µm. Resultant glass powders were used to fabricate glass ionomer cements with the addition of polyacrylic acid (PAA) and DI water. The exact formulation of the GICs is presented in Table 2. The initial characterization on GICs was to perform rheological testing of working and setting times. The setting time (St) of these cements is determined according to ISO 9917 standard, however the working time (Wt) is not governed by any standard so the Wt was taken as the period of time where the cement retains sufficient viscosity for implantation. The Wt and St were performed on the GICs made from 50wt% PAA concentration and is represented in seconds in Figure 1. From Figure 1 it can be seen that there is a compositional dependence in the Wt and St of each of these cements with an increased SrO concentration in the glass. The longest Wt of the cement series was attributed to GS0 with a Wt of 64s while the shortest Wt was with GS10 at with a Wt of 50s. This may be attributed to the Sr ion affecting the pH of the local environment of the cement mixture, hence resulting in a more rapid set. The setting times (St) are determined in accordance to the standards defined for the setting times of dental GICs. Figure 1 shows the St of GS0, GS5, and GS10 at 50wt% PAA concentration. The St of the cement series shown in Figure 1 behaves in a similar fashion to the Wt. The longest St of the cement series was attributed to GS0 with a St of 452s, which corresponds to 7 minutes and 32 seconds. The shortest St was attributed to GC10 with a time of 356s which corresponds to 5 minutes and 56 seconds.

**Figure 1.**
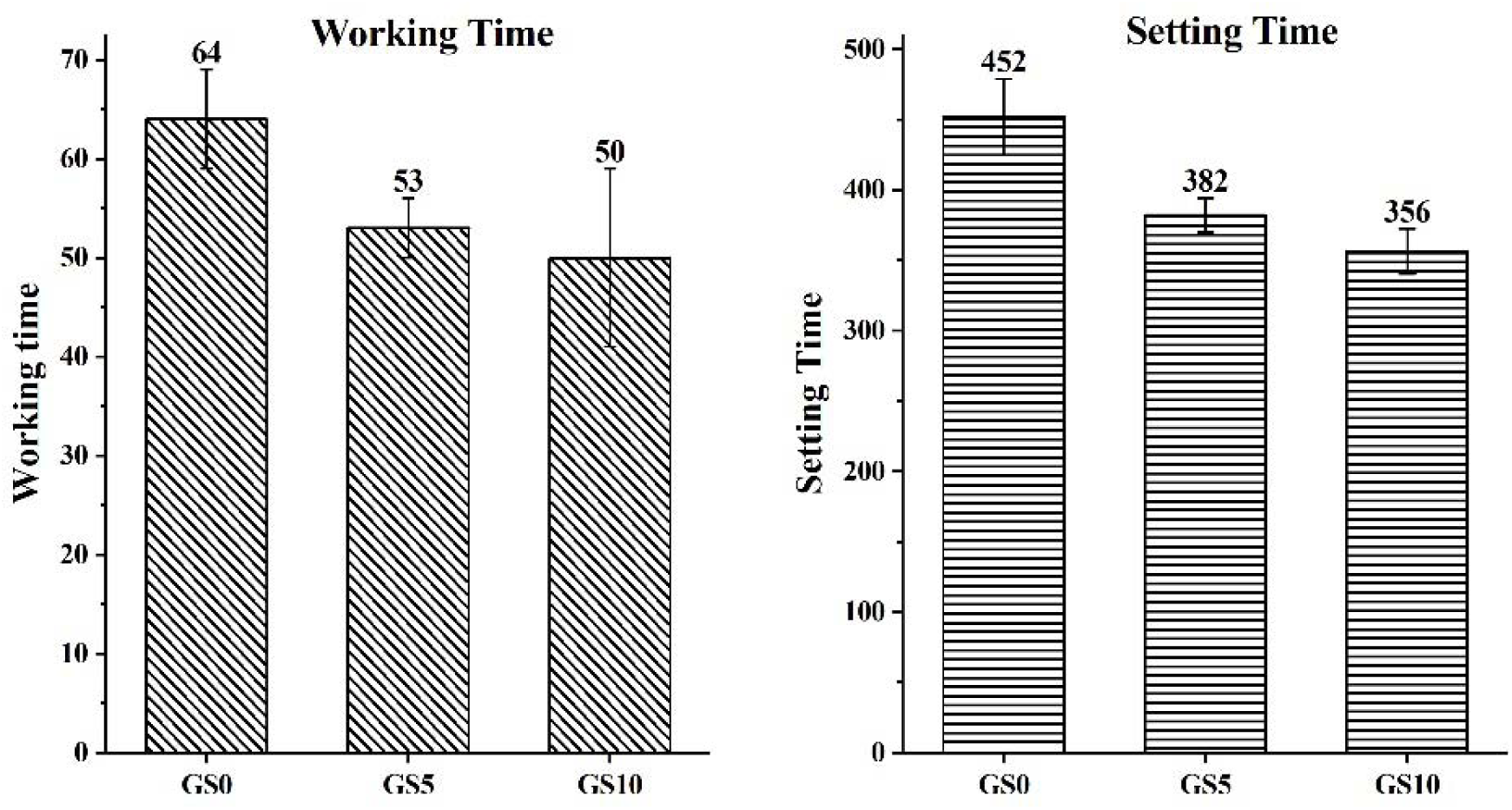
Working and Setting times of GS0, GS5, and GS10 cements.

An ideal bone cement needs to have a prolonged Wt and should set rapidly. The Wt allows the surgeon/user time to sufficiently mix the material to a homogenous paste and to properly manipulate the material into place. The bone cement should ideally then set rapidly. A rapid set cement reduces open wound exposure during surgery. Dental GICs are structurally more closely related to the cements discussed here than the orthopaedic bone cements and have been quoted in the literature as having St between 2.25 – 6.5 minutes depending on the composition R [1]. Cement series analyzed here exhibited St ranging between 7.32 to 5.56 minutes, which is close to the values reported for GICs. The Wt and St decreased by increasing the SrO content in the glass series. A possible explanation for the reduction in Wt and St could be due to the composition of the glass under investigation. Degradation of the glass toward the polyacrylic acid depends on the reactivity of the glass which is a great function of the glass chemistry and structure. Substitution of SiO_2_ for network modifiers, in this case SrO, will disrupt the glass network and make it susceptible to acid attack[20]. Therefore, the SrO substituted glasses will have faster degradation rates, and hence higher Wt and St compared to control glass with higher SiO_2_ content.

To analyze the mechanical properties of the cements, compressive strength and biaxial flexural strength of the cements were analyzed over 1, 10 and 30 days incubation in simulated body fluids (SBF). Figure 2 shows the compressive strength of the cements in MPa. After 1 day of incubation in SBF, the compressive strength of GS0, GS5, and GS10 was found to be at 21, 25, and 28 MPa, respectively. From Figure 2, it is evident that the compressive strength of the cements increases as a function of incubation time in the SBF. The highest compressive strength of all three compositions was found after 30 days of incubation period. The compressive strength of the cements after 30 days of incubation time was found to be at 27, 33, and 34 MPa for GS0, GS5, and GS10, respectively. Figure 2 also demonstrates that there is a compositional dependence in the compressive strength of the cements at each time period, where GS0 shows the lowest and GS10 shows the highest strength respectively.

**Figure 2.**
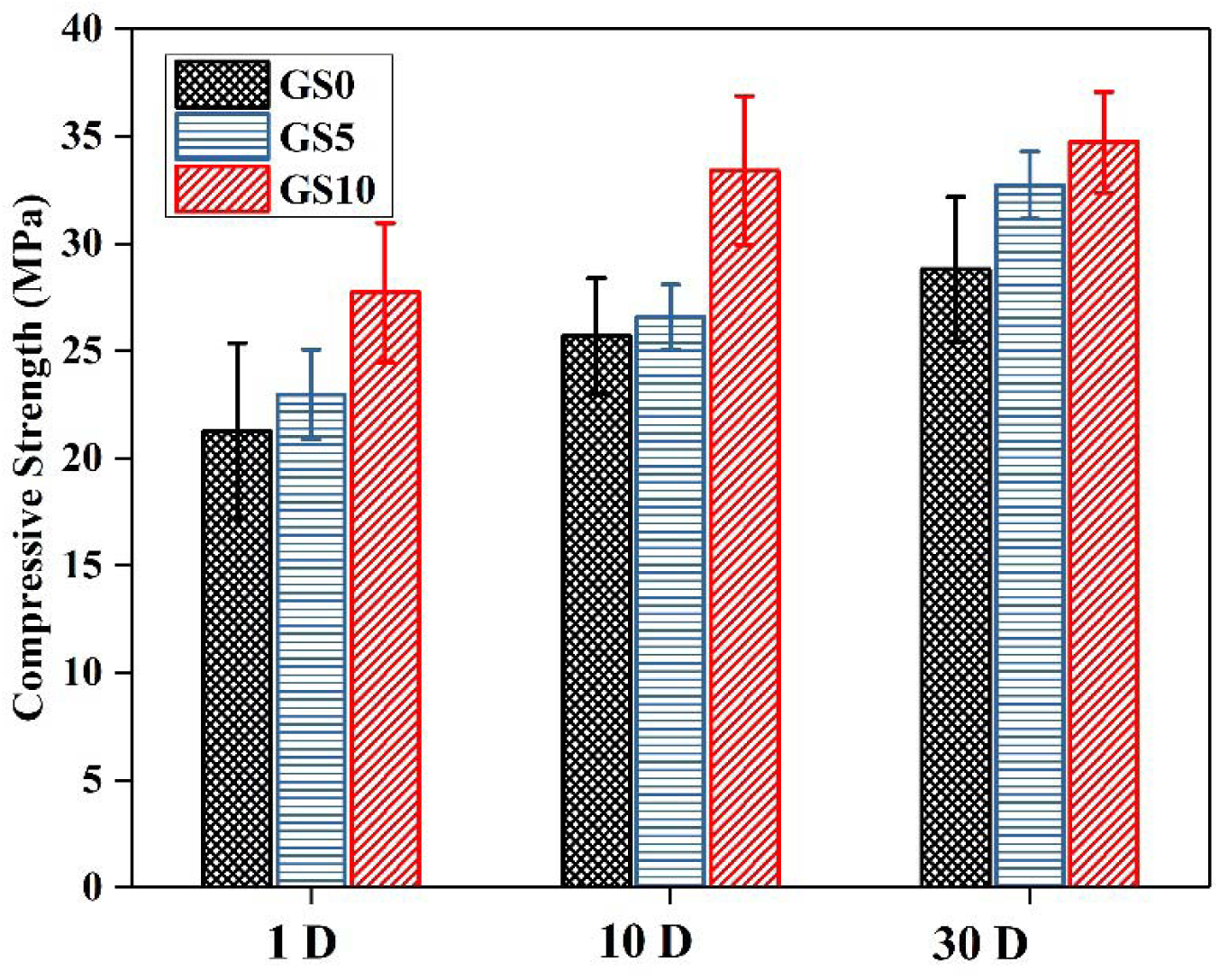
Compressive strength of GS0, GS5, and GS10 after 1, 10, and 30 days incubation in SBF.

The strengthening process in GICs is known to occur due to a number of parameters. The Mw and concentration of PAA used can influence strength. This occurs due to the increase in COO- groups which facilitates an increase in crosslinking within the cement matrix between the PAA chain and the metal cations released from the glass[25]. There is a compositional difference between the control and SrO containing cements. The Sr^2+^ in the glass phase of these cements acts as a network modifier where it facilitates the degree of crosslinking as compared to control cements. This is in agreement with the rheological studies of these cements with faster rates of Wt and St of SrO containing cements. Higher degradation rates in the SrO containing glasses and the release of Sr ions in the water within the process of setting of the cements, increases the rate of gelation. Unwinding of the PAA chains, and increased concentration of metallic cations in the solution can lead to an increas crosslinking density between the PAA chains and released ion within the cement matrix, which subsequently can increase the strength of the cements. Similarly, the increase in the strength as a function of time can be explained by the process of maturation of these cements which is associated with an increase in the crosslinking density. It is reported that the setting reaction of the commercial GICs, which is considered as dissolution and then gelation, take place within the first 24 hours[26]. After the initial setting reaction and over time, cements start to mature which is considered as the third stage of setting. Over maturation of the cements additional crosslinking occurs where it can potentially lead to increase in the mechanical properties. As cements mature, the steady release of ions from the glass phase into the solution will crosslink with the remaining unreacted PAA chain and facilitate the formation of new calcium, strontium polyacrylates which increases the crosslinking density. To further analyze the mechanical properties, biaxial flexural strength studies were performed on the cement series.

Figure 3 shows the biaxial flexural strength of the cement series as incubated in the SBF for 1, 10, and 30 days. From Figure 3, it is evident that the incubation of the cements did not significantly change the flexural strength of the cements. This is opposite to the trend observed for the compressive strength. The flexural strength of the GS0 was at 14.7, and 15.2 MPa after 1 and 30 days incubation time period, respectively. Similarly, for GS5 and GS10, the flexural strength was at 13.4 and 16.1 MPa after 1 day, and 14.7 and 16.3 MPa after 30 days of incubation, respectively. The results from Figure 3 indicate that the cements did not significantly change in the flexural strength during maturation process. The flexural strength test was originally developed for brittle ceramics however it is also suitable for brittle dental materials such as GICs. This material property tests the area of maximum tensile stresses which is located at the center of the lower face of the specimen. This is highly relevant as this test modality more accurately reflects the stresses present in a clinical situation, as bone typically fails in tension. Higgs et al also suggest that the measurement of strength in brittle materials under biaxial, rather than uniaxial conditions are often considered more reliable since edge failures (due to cement preparation) are eliminated[27]. The chemistry of the cements highly affects the compressive strength of the cements due to changes in the crosslinking density. However, in testing the flexural strength, internal defects such as voids and pores will significantly alter the flexural strength of the cements. Overall the mechanical properties testing of the cements revealed that they have sufficient mechanical properties to be considered for dental and skeletal applications. In particular, the compressive strength of the cements increased with maturation time, with SrO containing cements having higher values compared to the control cement.

**Figure 3.**
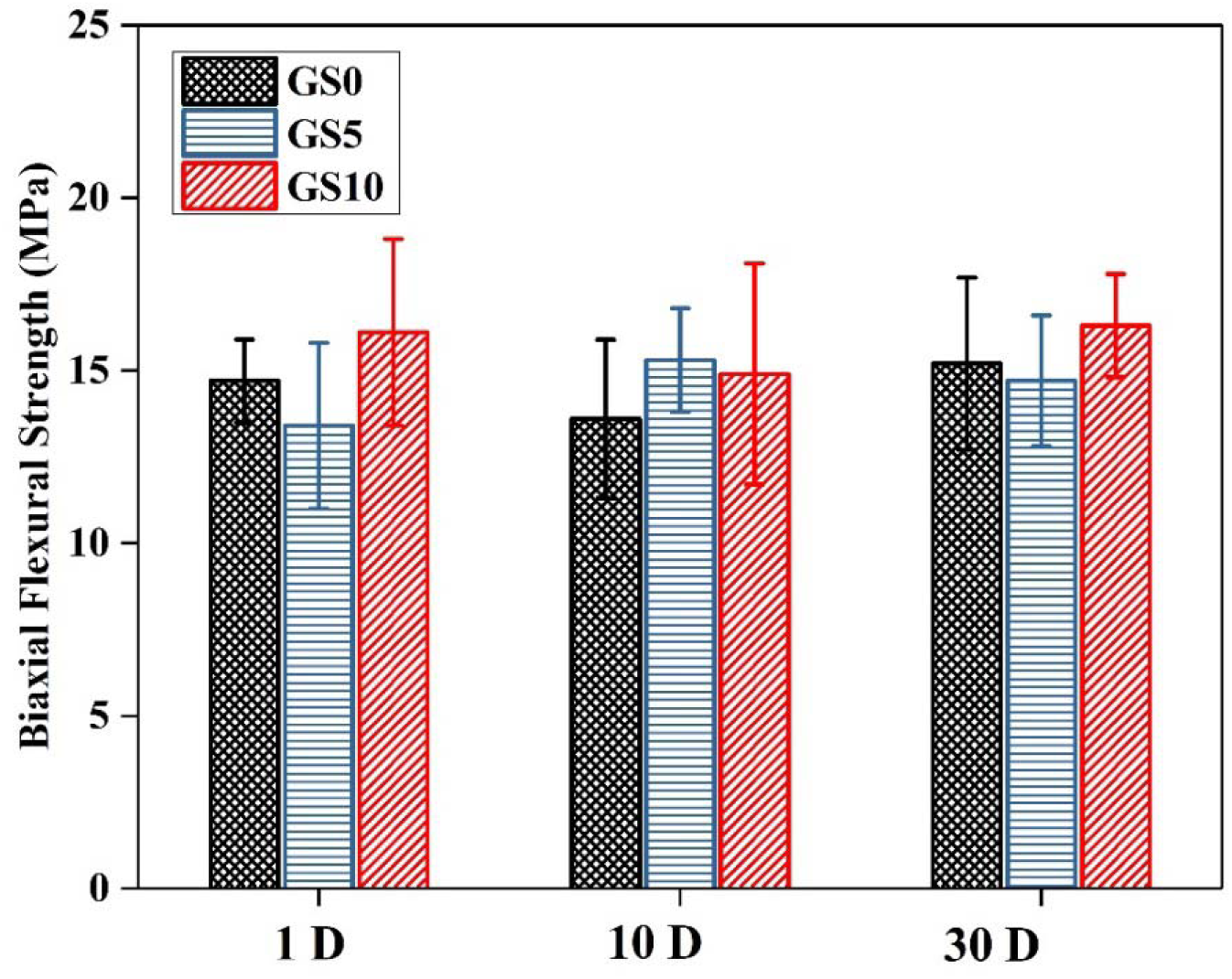
Biaxial Flexural strength of GS0, GS5, and GS10 after 1, 10, and 30 days incubation in SBF.

Another important characteristic that determines the lifespan of a biomaterial is the performance in the biological environment. A test that is widely accepted to determine a material’s performance in close proximity to bone is the Simulated Body Fluid (SBF) trials. The formation of calcium phosphate (CaP) layer on the surface, when immersed in SBF, is a good indication of its bioactivity. The deposition of CaP layer on the surface facilitates the nucleation of biological apatite. This layer then promotes adsorption of proteins, cell attachment and eventually the formation of a strong bond with the hard tissue[28]. SBF trials were conducted on the series of cements and the results are presented and discussed further. For SBF solution preparation, the method proposed by Kokubo et al was followed[24]. Figure 4 shows the SEM micrographs of the cement series incubated in SBF for 1 day. As can be seen from Figure 4, none of the cement series exhibited surface depositions after 1 day. These images show the presence of unreacted glass particles embedded in the polymer matrix. Cracks in the surface of the cement are caused by dehydration; a result of preparing these samples for SEM. The EDX trace for the control sample was found to contain only the reagents used to make the cement sample and is being used as a baseline to quantify how much calcium and phosphorus are present in the cement in comparison to the surface composition after immersion in SBF. Figure 5 shows the surface of the SBF incubated samples after 10 days. Control cement showed the least surface depositions at this time period whereas GS10 represented the highest amounts of the percipients. The presence of depositions on the surface of GS5, and GS10 samples can be attributed to CaP depositions. It is evident that CaP clusters are forming although they are widely dispersed. The elemental compositions of these precipitates were further analyzed by EDX. The surface depositions are increasing with respect to maturation time. After 30 days of incubation, surface of the all cement series is covered with the CaP precipitates as shown in Figure 6. The elemental compositions of the precipitates were analyzed using EDX, and results are presented in Figure 7. The EDX traces reveals the cement base materials Si, Zn, Sr and Ca of the cements along with high amount of P, Ca, and Zn at 23.1, 12.4, and 49.7 wt%.

**Figure 4.**
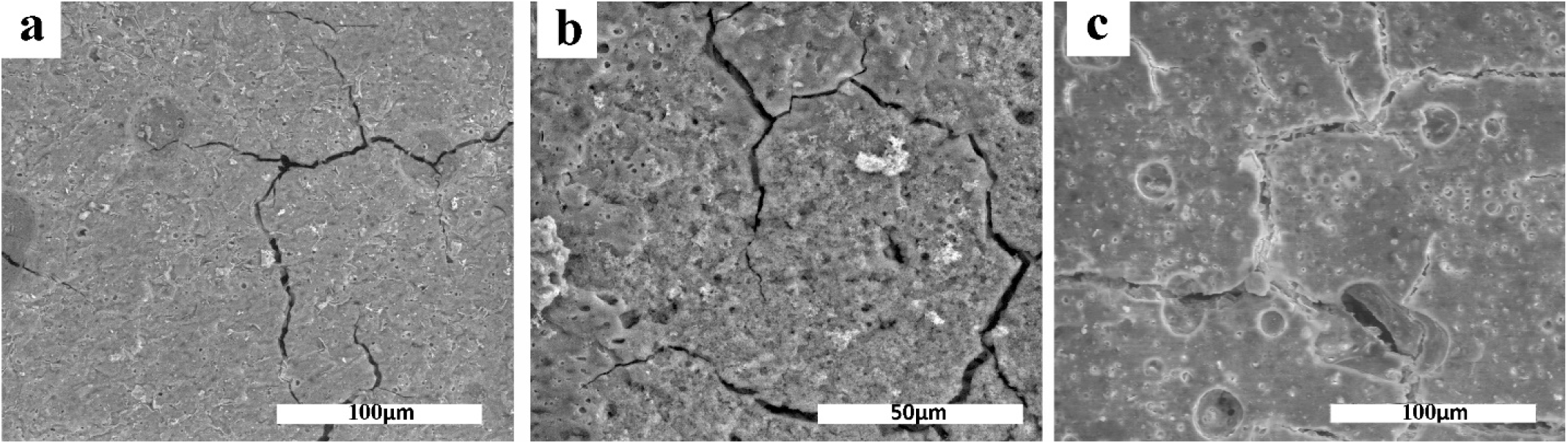
SEM images of a) GS0, b) GS5, and c) GS10 after 1-day incubation in SBF.

**Figure 5.**
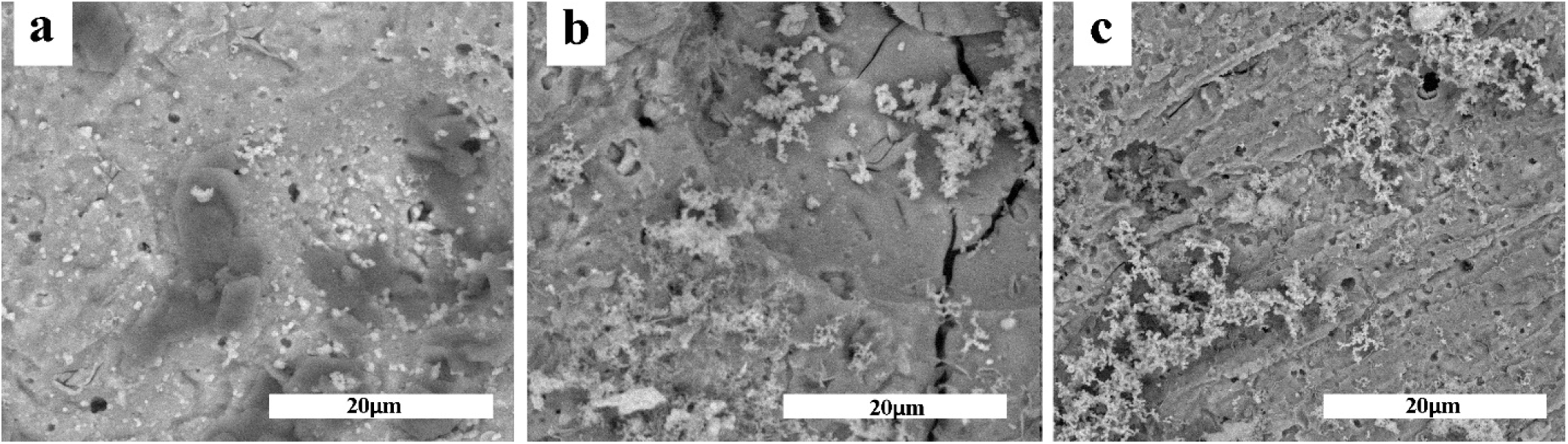
SEM images of a) GS0, b) GS5, and c) GS10 after 10-days incubation in SBF.

**Figure 6.**
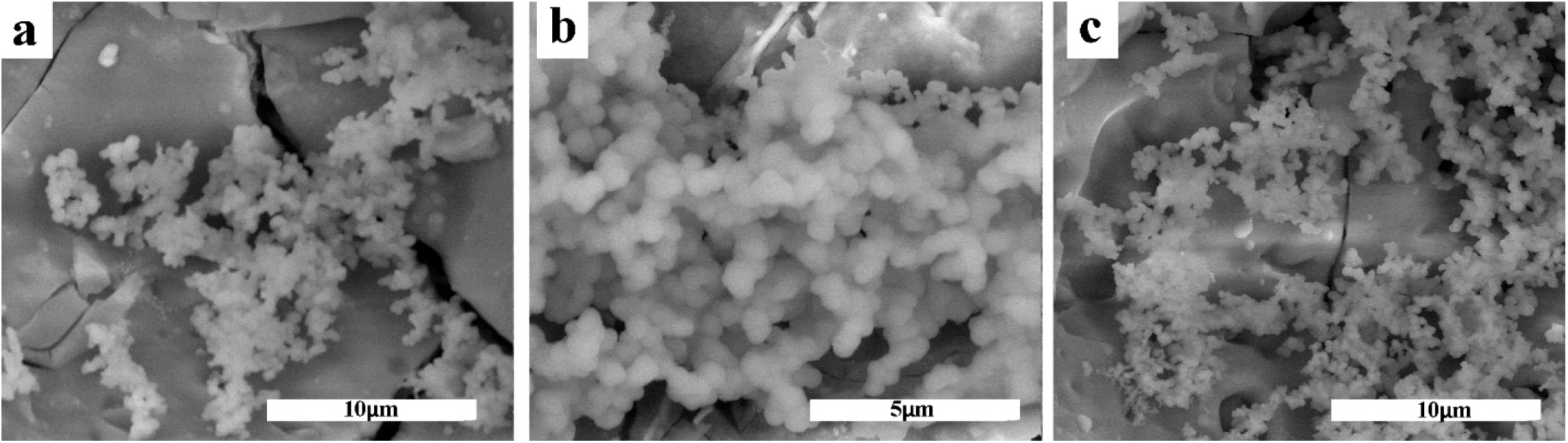
SEM images of a) GS0, b) GS5, and c) GS10 after 30-days incubation in SBF.

**Figure 7.**
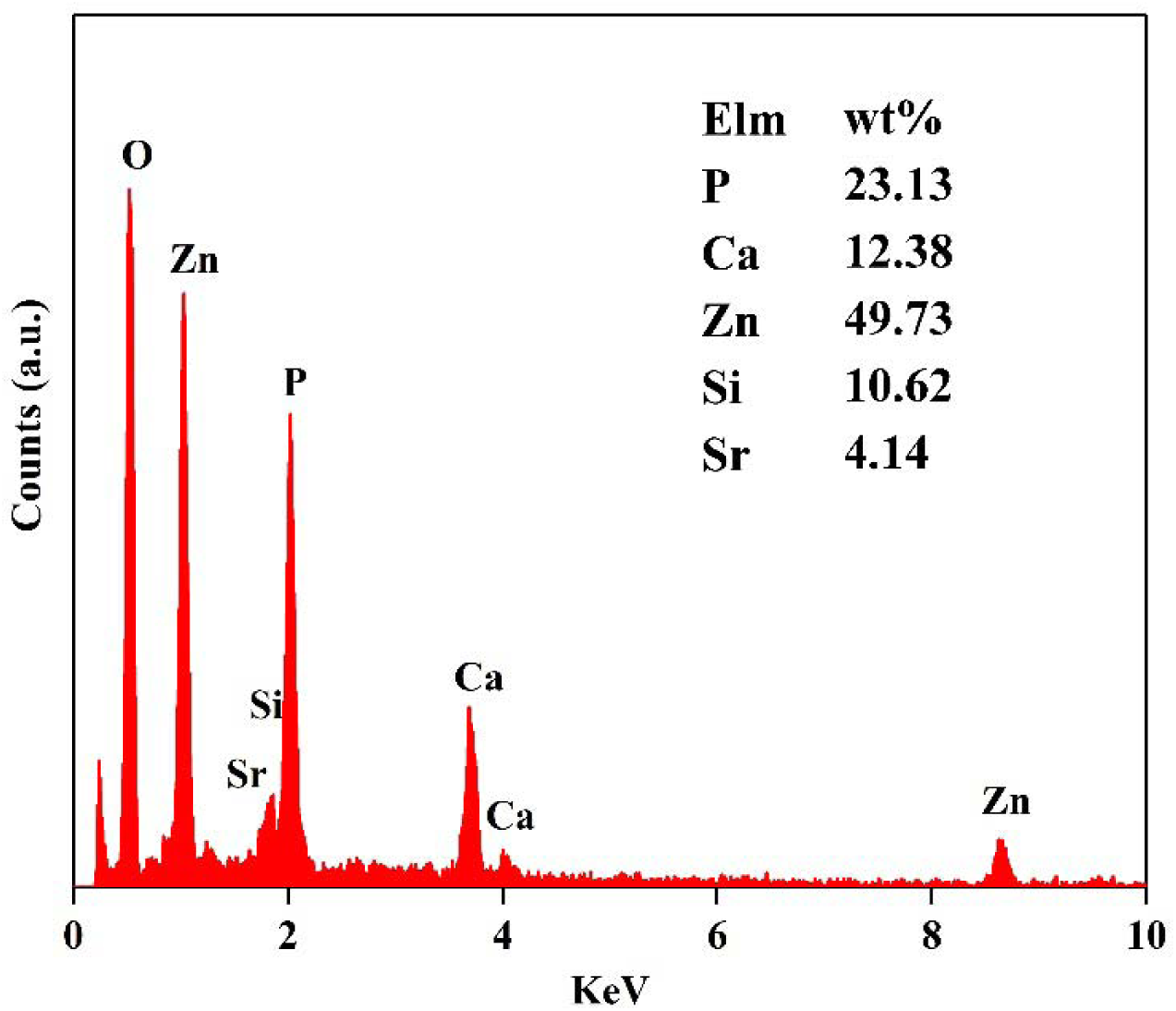
EDX analysis of the surface depositions on GS10 after 30 days incubation in SBF.

The *In-Vitro* bioactivity studies of novel GICs tested here show positive results when immersed in SBF. In particular, strontium containing cements, exhibited a higher amounts of CaP surface layer even after 10 days as identified by SEM. The main reason for the bioactivity of these GICs is the structure of the glass. It is known that precipitation of CaP is due to ion exchange between the cement and the surrounding solution[29]. The presence of network modifiers favors the ion exchange process by making the glass network disrupted. The incorporation of modifier ions in the silica matrix leads to a disruption of the glass network and formation of the non-bridging oxygen groups[29]. An increase in the number of non-bridging oxygen groups in the glass structure The disruption in the glass structure leads to a higher dissolution rate of the glass particles in the physiological solutions, which facilities the migration of Ca^2+^ and PO_4_^3-^ groups to the surface in order to form a CaP rich layer.

## 4 Conclusions

The substitution of SrO for SiO_2_ in the glass composition of the GICs was investigated in order to study the physical, mechanical and biological behavior of the resultant cements. Results from rheological behavior analysis indicated that the strontium incorporation reduced the working and the setting times of the cements. The addition of strontium resulted in a higher compressive strength of the cements as a result of higher crosslinking density. Bioactivity of the cements was analyzed using the SBF trials and after 1, 10, and 30 days. The presence of the CaP depositions was identified by SEM and EDX analysis. The positive results from the mechanical and biological behavior analysis of the strontium modified cements indicate their potential applications for skeletal tissue.

